# Serotonin inhibits axonal regeneration of identifiable descending neurons after a complete spinal cord injury in lampreys

**DOI:** 10.1101/335844

**Authors:** Daniel Sobrido-Cameán, Diego Robledo, Laura Sánchez, María Celina Rodicio, Antón Barreiro-Iglesias

**Affiliations:** Department of Functional Biology, CIBUS, Faculty of Biology, Universidade de Santiago de Compostela, 15782 Santiago de Compostela, Spain; The Roslin Institute and Royal (Dick) School of Veterinary Studies, The University of Edinburgh, Midlothian, EH25 9RG, UK; Department of Genetics, University of Santiago de Compostela, Campus de Lugo, Lugo, Spain

**Author notes:** Corresponding author: Dr. Antón Barreiro-Iglesias Address: Departamento de Biología Funcional, Edificio CIBUS, Campus Vida, Universidade de Santiago de Compostela, CP. 15782, Santiago de Compostela, A Coruña, Spain. Equal contributors (María Celina Rodicio, Antón Barreiro-Iglesias).

**Keywords:** Serotonin, serotonin receptor 1A, cAMP, spinal cord injury, lampreys, axon regeneration, morpholino, axonal guidance, Plexin-B1, ROBO

## Abstract

Classical neurotransmitters are mainly known for their roles as neuromodulators, but they also play important roles in the control of developmental and regenerative processes. Here, we used the lamprey model of spinal cord injury to study the effect of serotonin in axon regeneration at the level of individually identifiable descending neurons. Pharmacological and genetic treatments after a complete spinal cord injury showed that endogenous serotonin inhibits axonal regeneration in identifiable descending neurons through the activation of serotonin 1A receptors and a subsequent decrease in cAMP levels. RNA sequencing revealed that changes in the expression of genes that control axonal guidance could be a key factor on the serotonin effects during regeneration. This study provides new targets of interest for research in non-regenerating mammalian models of traumatic CNS injuries and extends the known roles of serotonin signalling during neuronal regeneration.

## Introduction

In contrast to mammals, including humans, lampreys recover locomotion spontaneously following a complete spinal cord injury (SCI; see ^1–4^). In lampreys, the process of recovery from SCI involves a positive astroglial response^5^, the production of new neurons in the spinal cord^6^ and the regeneration of ascending and descending axons through the injury site^7–10^. This regeneration is specific in that axons grow selectively in their normal directions^7,11,12^. Moreover, regenerated axons of descending brain neurons are able to re-establish synaptic connections with their appropriate targets below the site of injury^13,14^. However, even in lampreys not all descending neurons of the brain are able to regenerate their axons following a complete spinal cord transection, even when normal appearing locomotor function is observed several weeks after the injury^8,9^. Approximately, 50% of all descending brainstem neurons are able to regenerate their axon below the site of injury after a complete SCI^14,15^. Among brain descending neurons, the brainstem of lampreys contains several giant individually identifiable descending neurons that vary greatly in their regenerative ability, even when their axons run in similar paths in a spinal cord that is permissive for axonal regrowth^9^. Some identifiable neurons like the Mauthner or I1 neurons regenerate their axon less than 10% of the times after a complete spinal cord transection, while other identifiable neurons like the I3 or B6 neurons are able to regenerate their axon more than 60% of the times after a complete SCI^9^. This suggests that intrinsic factors present in some neurons but not in others might limit their regenerative ability. Lampreys offer a convenient vertebrate model in which to study the inhibition or promotion of axonal regeneration after SCI in the same *in vivo* preparation.

One of the main molecules known to be an intrinsic regulator of axonal regeneration is the second messenger cyclic-adenosine monophosphate (cAMP) (see^16,17^). Several reports in mammals and fishes have shown that cAMP promotes axon regeneration following SCI^18–22^. Subsequent studies have also shown that cAMP promotes axon regeneration in descending neurons of lampreys after SCI^23–25^. The challenge now is to define the signals that control cAMP levels in descending neurons after axotomy and during regeneration.

Several neurotransmitters modulate intracellular cAMP levels by activating metabotropic G-protein coupled receptors, which include serotonin, glutamate, GABA or dopamine receptors. So, neurotransmitters acting through these receptors are potential regulators of intracellular cAMP levels following a traumatic injury to the CNS. Among them, serotonin appears as a good candidate to regulate axon regeneration following nervous system injuries (see^26^). Serotonin receptors are divided in 7 families, with families 1, 2 and 4 to 7 being G-protein coupled metabotropic receptors and type 3 serotonin receptors being ligand-gated ion channels. Families 1 and 5 of serotonin receptors are known to decrease intracellular levels of cAMP; whereas, families 4, 6 and 7 increase intracellular levels of cAMP. A few *in vitro* and *ex vivo* studies have shown that serotonin inhibits axon regrowth in invertebrate^27,28^ and vertebrate^29,30^ species. In contrast, a recent *in vivo* study in *Caenorhabditis elegans* showed that 5-HT promotes axon regeneration after axotomy^31^. However, no study has yet looked at the role of serotonin in axon regeneration in a *in vivo* vertebrate model of traumatic CNS injury. The presence of rich serotonin innervation in the vicinity of descending neurons of the lamprey brainstem^32,33^, the expression of serotonin 1A receptors in identifiable descending neurons of lampreys^34^ and data showing an increase in synaptic contacts on descending neurons following SCI in lampreys^35^ prompted us to study the possible role of serotonin in axon regeneration following SCI in lampreys. Here, we present gain and loss of function data using pharmacological and genetic treatments showing that endogenous serotonin inhibits axon regeneration in identifiable descending neurons of lampreys following a complete SCI by activating serotonin 1A receptors. We also performed an RNA sequencing study, which revealed that changes in the expression of genes that control axonal guidance could be a key factor on the serotonin effects during regeneration. This provides a new target of interest for SCI research in non-regenerating mammalian models.

## Results

### A serotonin treatment inhibits axon regeneration in identifiable descending neurons after a complete spinal cord injury

To reveal the effect of serotonin in the regeneration of identifiable descending neurons, larval sea lampreys were treated with the serotonin analogue serotonin-hydrochloride for a month following a complete spinal cord transection. At 11 weeks post-lesion (wpl), the serotonin treatment significantly inhibited axon regeneration of identifiable descending neurons of the sea lamprey (Student’s t-test, *p* = 0.0145; Fig. 1A-C). Interestingly, behavioural analyses revealed that the serotonin treatment did not cause a general toxic effect since locomotor recovery was not affected by the serotonin treatment (Fig. 1D).

**Figure 1.**
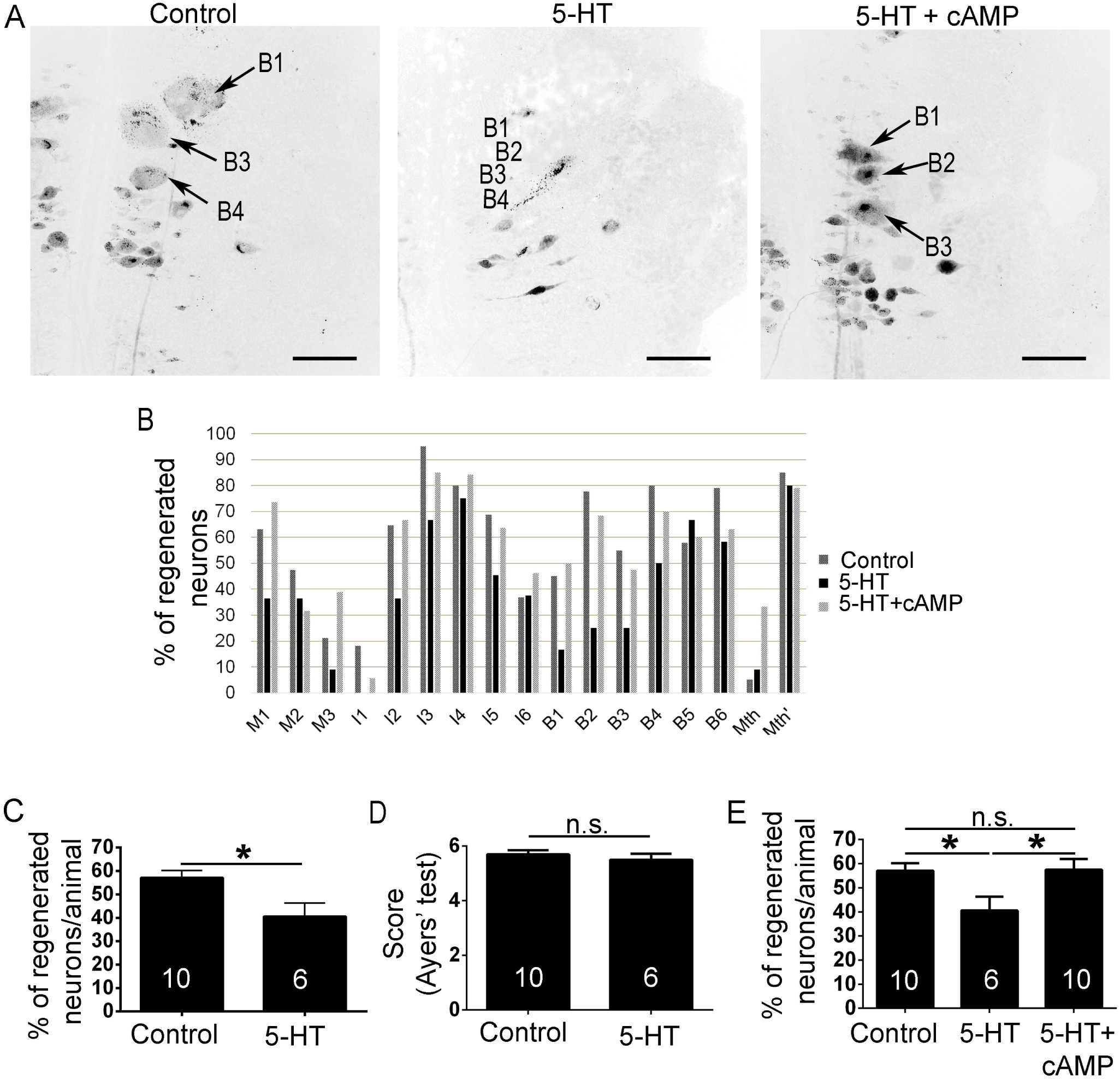
A serotonin (5-HT) treatment inhibits axonal regeneration of identifiable descending neurons and an additional cAMP treatment rescues the inhibitory effect of 5-HT. **A:** Photomicrographs of whole-mounted brains showing regenerated identifiable neurons, as identified by retrograde labelling, in control, 5-HT treated and 5-HT and cAMP treated animals. Note the decreased number of labelled (regenerated) identifiable neurons in 5-HT treated animals. **B:** Graph showing the percentage of regenerated neurons (respect to the total number of analysed neurons) for each identifiable descending neuron in control, 5-HT treated and 5-HT and cAMP treated animals. **C:** Graph showing significant changes (asterisk) in the percentage of regenerated neurons per animal after the 5-HT treatment (control: 57.06 ± 3.08 %; 5-HT treated: 40.56 ± 5.74 %). **D:** Graph showing non-significant differences in recovery of swimming activity (Ayers’ test) in control and 5-HT treated animals (Ayer’s test score: control: 5.70 ± 0.15; 5-HT treated: 5.50 ± 0.22). **E:** Graph showing significant changes (asterisks) in the percentage of identifiable regenerated neurons per animal after the 5-HT or 5-HT and cAMP treatments (control: 57.06±3.08 %; 5-HT: 40.56 ± 5.74 %; 5-HT + cAMP: 57.51 ± 4.39 %). Arrows indicate identifiable descending neurons that regenerated in control or 5-HT and cAMP treated animals, but not in 5-HT treated animals. Rostral is up and the midline to the left in all photomicrographs. Scale bars: 100 µm.

### cAMP treatment can reverse the inhibitory effect of serotonin on axonal regeneration

Previous studies have shown that a single dose of cAMP applied at the time of transection promotes axon regeneration of identifiable descending neurons following a complete spinal cord transection in larval sea lampreys^23–25^. Here, and to reveal a possible involvement of this second messenger in the inhibitory effect of serotonin in axonal regeneration, we carried out a rescue experiment in which animals treated with serotonin as above were also treated with dibutyryl-cAMP (db-cAMP). The db-cAMP treatment was able to significantly rescue the inhibitory effect of serotonin, with these animals reaching levels of axonal regeneration of identifiable descending neurons similar to control vehicle treated animals (ANOVA, *p* = 0.0288, Fig. 1A, B and E).

### Correlation between the expression of 5-HT1A and the regenerative ability of identifiable neurons

The serotonin 1A receptor is an ancient G-protein coupled receptor that is known to decrease intracellular cAMP levels through Gi/Go when activated by serotonin. In a previous study, we showed that this receptor is expressed in identifiable descending neurons of larval sea lampreys^34^. This, together with the results of the db-cAMP treatments (see previous section) prompted us to study whether there was a correlation between the expression of the serotonin 1A receptor in identifiable descending neurons and their known regenerative ability following a complete spinal cord transection. First, qPCR analyses of the whole brainstem (where most descending neurons are located) revealed no significant changes in the expression of the serotonin 1A receptor 4 wpl as compared to control animals (Suppl. Fig. 1). We should take into account that the brainstem contains a large number of non-descending neurons and also that only about 50% of descending neurons of the lamprey brainstem regenerate after a complete spinal cord transection (see introduction). So, we decided to carry out *in situ* hybridization analyses in horizontal sections of the sea lamprey brain to look at the expression of the serotonin 1a receptor in individually identifiable neurons. This, as opposed to whole-mounts, impedes the clear identification of all identifiable descending neurons in all sections; therefore, in these analyses only the M1, M2, M3, I1 and I3 neurons where included.

Our *in situ* hybridization experiments revealed that in bad regenerator neurons (neurons that regenerate their axon less than 30% of the times) there is a significant increase in the expression of the serotonin 1A receptor 4 weeks after a complete SCI (M2 neuron: Kruskal-Wallis, *p* = 0.0105; M3 neuron: ANOVA, *p* = 0.0478; I1 neuron: ANOVA, *p* = 0.0238; Fig. 2A; Table 1), while in good regenerator neurons (M1 and I3 neurons) the expression of the receptor decreases (non-significantly) in the first weeks following a complete spinal cord transection (Fig. 2B; Table 1). Statistical analyses revealed a significant correlation between the percentage of increase/decrease in the expression of the receptor 4 wpl and the long-term regenerative ability of identifiable neurons (Pearson’s test, *p* = 0.0293, Fig. 2C). This suggests that the presence and activity of this receptor in descending neurons might inhibit axonal regeneration after a complete SCI.

**Table 1.** Mean ± S.E.M. values of the number of 5-ht1a positive *in situ* pixels per section in identifiable descending neurons of control and injured animals. Refers to Fig. 2.

| 5-HT1a positive<br>pixels/section | Control | 1 wpl | 4 wpl |
| --- | --- | --- | --- |
| M1 | 118,053±28,907 | 69,944±29,647 | 47,977±23,429 |
| M2 | 12,188±4,609 | 22,354±11,186 | 73,734±28,394 |
| M3 | 17,981±7,496 | 74,329±28,886 | 136,996±52,416 |
| I1 | 9,367±5,697 | 15,864±10,099 | 19,685±8,690 |
| I3 | 52,673±20,947 | 7,698±3,587 | 19,714±10,667 |

**Figure 2.**
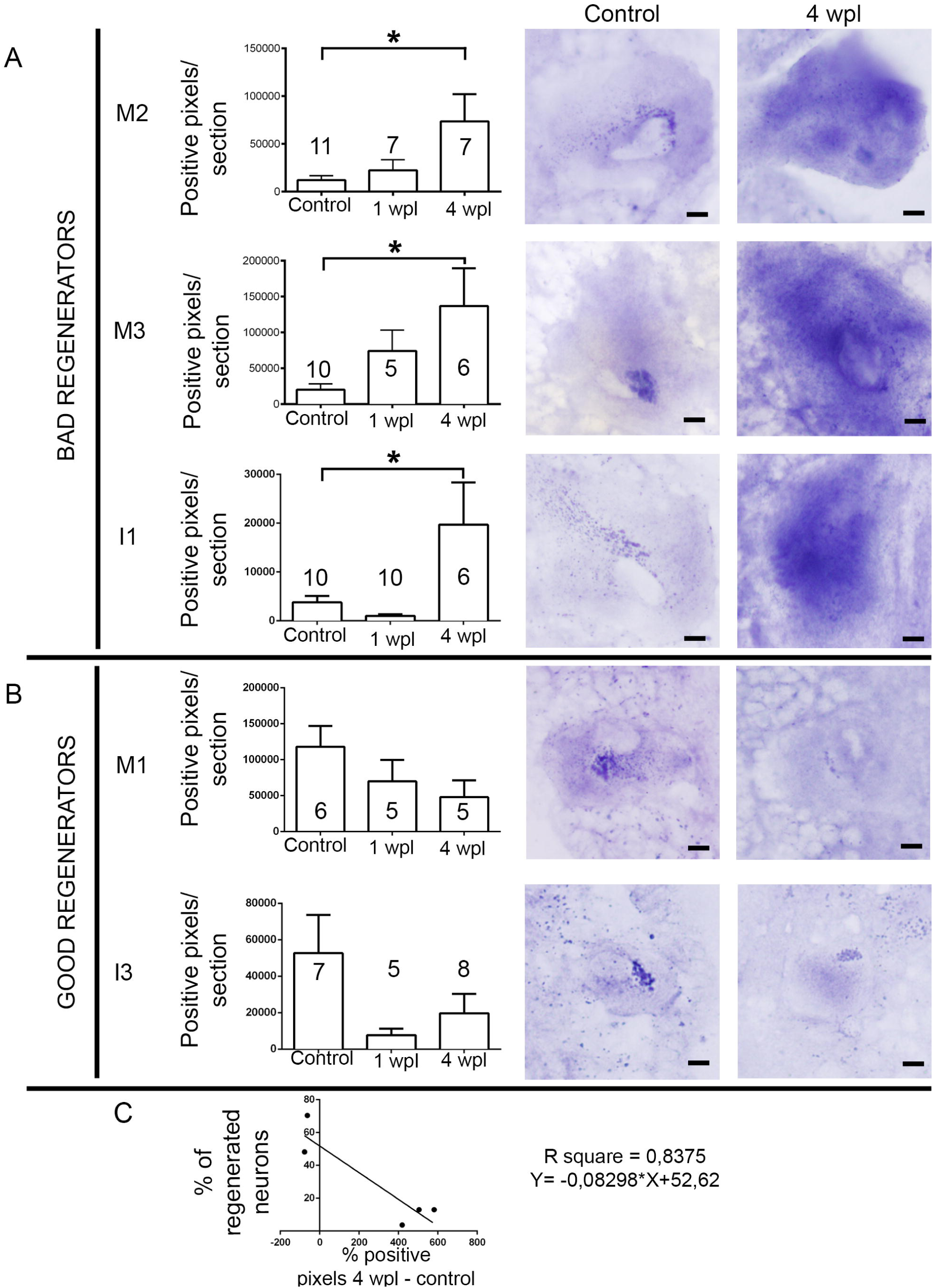
Changes in the expression of the serotonin 1a receptor (5-ht1a) in identifiable descending neurons after a complete SCI. **A:** Graphs and photomicrographs showing significant changes (asterisk) in the number of 5-ht1a positive *in situ* pixels per section of the soma of bad regenerator identifiable descending neurons. The mean ±S.E.M. values are provided in table 2. **B:** Graphs and photomicrographs showing changes in the number of 5-ht1a positive *in situ* pixels per section of the soma of good regenerator identifiable descending neurons. The mean ± S.E.M. values are provided in table 2. **C:** Linear regression analysis showed an inverse relation between the percentage of change in 5ht1a transcript expression at 4 wpl and regeneration probability (from Jacobs et al., 1997) of each cell (95% confidence intervals for slope =-0.08298 ± 0.0211; Sy.x = 13.23). Scale bars: 20 µm.

### Endogenous serotonin inhibits axon regeneration through the 5-HT1A receptor following SCI

Based on the *in situ* hybridization results, we decided to study whether endogenous serotonin inhibits axon regeneration of identifiable descending neurons by activating serotonin 1A receptors. For this, we treated animals after a complete spinal cord transection with the serotonin 1A receptor antagonist WAY-100,135. Spinal cord transected animals were treated with WAY-100,135 for 4 weeks following a complete spinal cord transection. The WAY-100,135 treatment significantly promoted axon regeneration 11 wpl in identifiable neurons compared to control vehicle treated animals (Student’s t-test, *p* = 0.049, Fig. 3A-C). This indicates that endogenous serotonin inhibits axon regeneration following a complete SCI by activating serotonin 1A receptors.

**Figure 3.**
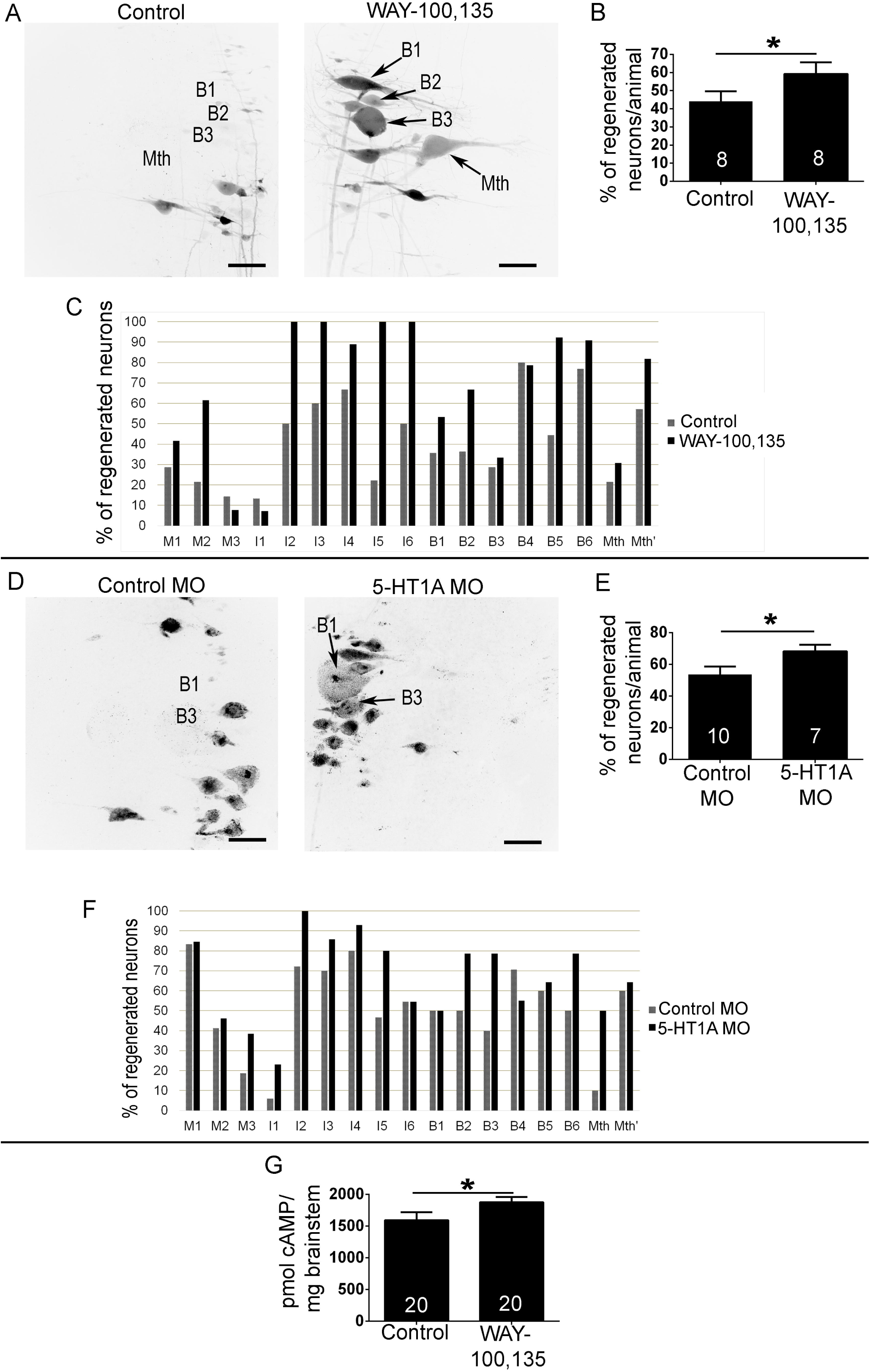
WAY-100,135 or 5-HT1A MO treatments promote axonal regeneration of identifiable descending neurons. **A:** Photomicrographs of whole-mounted brains showing regenerated identifiable neurons, as identified by retrograde labelling, in control and WAY-100,135 treated animals. Note the increased number of labelled (regenerated) identifiable neurons in WAY-100,135 treated animals. **B:** Graph showing significant changes (asterisk) in the percentage of regenerated neurons per animal after the WAY-100,135 treatment (control: 44.05 ± 5.58 %; WAY-100,135: 59.16 ± 6.44 %). **C:** Graph showing the percentage of regenerated neurons (respect to the total number of analysed neurons) for each identifiable cell in control and WAY-100,135 treated animals. **D:** Photomicrographs of whole-mounted brains showing regenerated identifiable neurons, as identified by retrograde labelling, in control and 5-HT1A MO treated animals. Note the increased number of labelled (regenerated) identifiable neurons in 5-HT1A MO treated animals. **E:** Graph showing significant changes (asterisk) in the percentage of regenerated neurons per animal after the 5-HT1A MO treatment (control: 53.69 ±4.97 %; 5-HT1A MO: 68.29 ± 4.12 %). **F:** Graph showing the percentage of regenerated neurons (respect to the total number of analysed neurons) for each identifiable cell in control and 5-HT1A MO treated animals. **G:** Graph showing significant changes (asterisk) in cAMP concentration per milligram of brainstem in control and WAY-100,135 treated animals (control: 1.594 ± 125.4 pmol of cAMP/mg of brainstem; WAY-100,135: 1.878 ± 81.74 pmol of cAMP/mg of brainstem). Arrows indicate descending neurons that regenerated in WAY-100,135 or 5-HT1A MO treated animals but not in controls animals. Rostral is up in all photomicrographs. The midline is to the right in photomicrographs of control animals and to the left in photomicrographs of treated animals. Scale bars: 100 µm.

**Figure 4.**
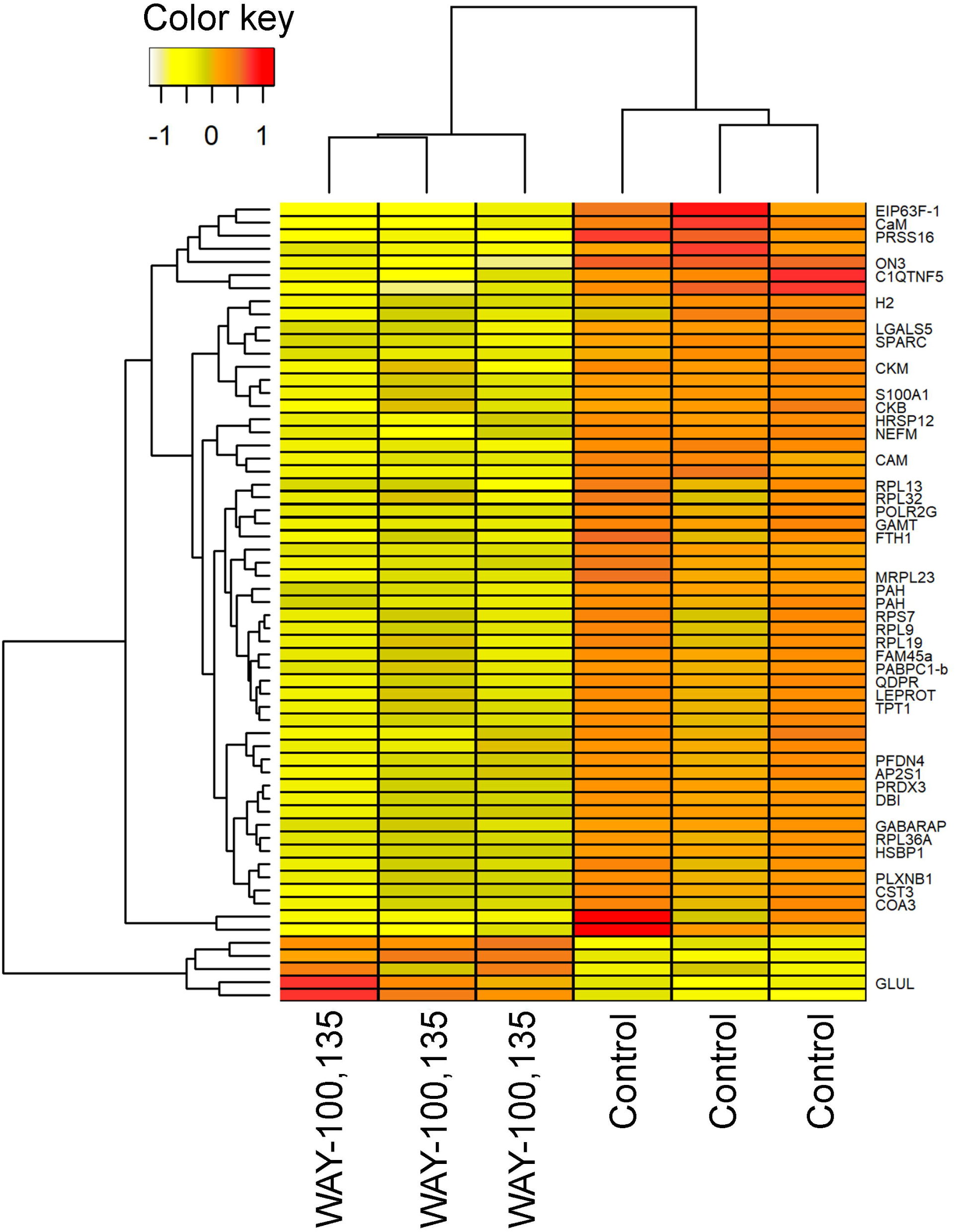
Heatmap showing the expression of 61 differentially expressed genes between samples treated with WAY-100,135 and control samples in RNA sequencing. Gene expression is represented as regularized log-transformed read counts scaled by gene. Samples (x-axis) and genes (y-axis) were hierarchically clustered according to their gene expression Euclidian distances using complete linkage clustering.

To confirm that the inhibitory effect of endogenous serotonin is due to the activation of serotonin 1A receptors expressed in identifiable descending neurons we decided to specifically knock-down the expression of the receptor in these neurons by using morpholinos targeted against the translation initiation site of the 5-HT1A receptor sequence (Suppl. Fig. 2A). The morpholinos were applied at the time of transection on the rostral stump of the spinal cord. Fluorescent labelling of the morpholinos confirmed that these were retrogradely transported and reached the soma of descending neurons during the first wpl (Suppl. Fig. 2B). Immunohistochemical analyses revealed that at 4 wpl the application of the active morpholino significantly reduced the expression of the serotonin 1A receptor in reticulospinal neurons of the sea lamprey brainstem as compared to the standard control morpholino (Student’s t-test, *p* = 0.001; Suppl. Fig. 2C, D). More importantly, and as expected from the antagonist treatments, the application of the 5-HT1A morpholino significantly promoted axon regeneration of identifiable neurons following a complete spinal cord transection as compared to the animals that received the standard control morpholino (Mann Whitney U test, *p* = 0.0177, Fig. 3D-F). Both the antagonist and morpholino treatments indicate that endogenous serotonin inhibits axon regeneration in identifiable neurons after SCI by activating serotonin 1A receptors expressed in these neurons.

### The WAY-100,135 treatment increases cAMP levels in the brainstem

To study whether changes in cAMP levels might be behind the beneficial effects of the serotonin 1A receptor antagonist treatments another set of animals was treated with WAY-100,135 for 1 week. A cAMP detection assay of the whole-brainstem revealed that the antagonist treatment, which also promoted axonal regeneration (see previous section), significantly increased cAMP levels in the brainstem (Mann-Whitney U test, *p* = 0.0397; Fig. 3G). This result indicates that the inhibition of axon regeneration caused by the activation of serotonin 1A receptors by endogenous serotonin might be caused by a decrease in intracellular levels of cAMP.

### RNA sequencing following WAY-100,135 treatment reveals new signalling pathways possibly involved in axonal regeneration in lampreys

Our gain and loss of function experiments revealed that endogenous serotonin inhibits axon regeneration following SCI in the sea lamprey through the activation of serotonin 1A receptors and that this effect might be caused by a decrease in intracellular cAMP levels. To reveal new genes that might be involved in the intrinsic control of axon regeneration and whose expression is modulated by the activity of 1A receptors we decided to repeat the WAY-100,135 treatment and carry out an RNA sequencing study in the sea lamprey brainstem at 4 wpl.

A total of 61 differentially expressed genes were detected between WAY-100,135 treated samples and control vehicle treated animals (Figure 4, Supplementary Table 1). Most of these genes were found down-regulated after WAY-100,135 treatment (5 up-regulated genes, 56 down-regulated). Among these down-regulated genes in response to WAY-100,135 is Plexin-B1 (PLXNB1, logFC = −0.42), which plays a role in axon guidance and works as a transmembrane receptor for semaphorins^36^. Only one of the five significantly up-regulated genes was annotated, Glutamine synthetase (GLUL, logFC = 1.05). This gene is primarily found in astrocytes and protects neurons against excitotoxicity^37^. Pathway analysis using Reactome^38^ revealed 29 significantly enriched pathways (FDR p-value < 0.05; Supplementary Table 2). Among these, the most interesting ones are “Axon guidance”, “Signalling by ROBO receptors” and “Regulation of expression of SLITs and ROBOs”, which are represented only by down-regulated genes.

**Table 2.** Table showing the number of animals included in each experimental group and also the total number of identifiable descending neurons that were included in the analyses. Note that in the *in situ* hybridization experiments only the neurons that were unequivocally identified in at least 2 brain sections were included in the quantifications.

|  |  | Total number of neurons included in the analyses |  |  |  |  |  |  |  |  |  |  |  |  |  |  |  |  |  |
| --- | --- | --- | --- | --- | --- | --- | --- | --- | --- | --- | --- | --- | --- | --- | --- | --- | --- | --- | --- |
|  |  | Animals | M1 | M2 | M3 | I1 | I2 | I3 | I4 | I5 | I6 | B1 | B2 | B3 | B4 | B5 | B6 | Mth | Mth' |
| 5-HT + cAMP<br>treatments<br>(11 wpl) | Control | 10 | 19 | 19 | 19 | 11 | 17 | 20 | 20 | 16 | 19 | 20 | 18 | 20 | 20 | 19 | 19 | 20 | 20 |
|  | 5-HT<br>treated | 6 | 11 | 11 | 11 | 11 | 11 | 12 | 12 | 11 | 8 | 12 | 12 | 12 | 12 | 12 | 12 | 11 | 10 |
|  | 5-HT +<br>cAMP treated | 10 | 19 | 19 | 18 | 18 | 18 | 20 | 19 | 11 | 13 | 20 | 19 | 19 | 20 | 20 | 19 | 18 | 19 |
| **5-ht1a qPCR** | Control | 8 | /// | /// | /// | /// | /// | /// | /// | /// | /// | /// | /// | /// | /// | /// | /// | /// | /// |
|  | 4 wpl | 8 | /// | /// | /// | /// | /// | /// | /// | /// | /// | /// | /// | /// | /// | /// | /// | /// | /// |
| **5-ht1a *in situ* hybridization** | Control | 6 | 11 | 10 | 10 | 8 | /// | 7 | /// | /// | /// | /// | /// | /// | /// | /// | /// | /// | /// |
|  | 1 wpl | 5 | 7 | 5 | 10 | 8 | /// | 5 | /// | /// | /// | /// | /// | /// | /// | /// | /// | /// | /// |
|  | 4 wpl | 5 | 7 | 6 | 6 | 7 | /// | 6 | /// | /// | /// | /// | /// | /// | /// | /// | /// | /// | /// |
| **WAY-100,135 treatment (11 wpl)** | Control | 8 | 14 | 14 | 14 | 15 | 14 | 15 | 15 | 9 | 4 | 14 | 11 | 14 | 15 | 9 | 13 | 14 | 14 |
|  | Treated | 8 | 12 | 13 | 13 | 14 | 10 | 11 | 9 | 5 | 3 | 15 | 12 | 15 | 14 | 13 | 11 | 13 | 11 |
| **5-HT1A MO treatment (11 wpl)** | Control | 10 | 19 | 18 | 19 | 17 | 17 | 19 | 19 | 14 | 12 | 22 | 22 | 22 | 22 | 22 | 22 | 22 | 22 |
|  | Treated | 7 | 13 | 13 | 13 | 13 | 13 | 14 | 14 | 9 | 11 | 14 | 14 | 14 | 14 | 14 | 14 | 14 | 14 |
| **5-HT1A MO retrograde transport (1 wpl)** | Control | 2 | /// | /// | /// | /// | /// | /// | /// | /// | /// | /// | /// | /// | /// | /// | /// | /// | /// |
|  | Treated | 2 | /// | /// | /// | /// | /// | /// | /// | /// | /// | /// | /// | /// | /// | /// | /// | /// | /// |
| **5-HT1A immunohistochemistry (MO treated, 4 wpl)** | Control | 5 | /// | /// | /// | /// | /// | /// | /// | /// | /// | /// | /// | /// | /// | /// | /// | /// | /// |
|  | Treated | 6 | /// | /// | /// | /// | /// | /// | /// | /// | /// | /// | /// | /// | /// | /// | /// | /// | /// |
| **5-HT1A immunohistochemistry (Pre-adsorption)** | Control | 3 | /// | /// | /// | /// | /// | /// | /// | /// | /// | /// | /// | /// | /// | /// | /// | /// | /// |
|  | Pre-adsorpted | 3 | /// | /// | /// | /// | /// | /// | /// | /// | /// | /// | /// | /// | /// | /// | /// | /// | /// |
| **cAMP assay (1 wpl)** | Control | 20 | /// | /// | /// | /// | /// | /// | /// | /// | /// | /// | /// | /// | /// | /// | /// | /// | /// |
|  | Treated | 20 | /// | /// | /// | /// | /// | /// | /// | /// | /// | /// | /// | /// | /// | /// | /// | /// | /// |
| **RNA-seq (4 wpl)** | Control | 12 | /// | /// | /// | /// | /// | /// | /// | /// | /// | /// | /// | /// | /// | /// | /// | /// | /// |
|  | Treated | 12 | /// | /// | /// | /// | /// | /// | /// | /// | /// | /// | /// | /// | /// | /// | /// | /// | /// |
| **<u>TOTAL</u>** |  | **176** |  |  |  |  |  |  |  |  |  |  |  |  |  |  |  |  |  |

## Discussion

We have provided gain and loss of function evidence, using pharmacological and genetic treatments, showing that endogenous serotonin inhibits axon regeneration after SCI in lampreys by activating serotonin 1A receptors expressed in descending neurons. Our results also suggest that differential changes in serotonin 1A receptor expression in descending neurons could be one of the factors that explain the different regenerative abilities of individual descending neurons.

Only a few studies have previously looked at the possible role of serotonin in neurite/axon regeneration and these have been carried out only in invertebrate and *in vitro* models (see^26^). Koert and co-workers^28^ showed that in the snail *Lymnaea stagnalis* auto-released serotonin inhibited neurite outgrowth in cultured serotonergic cerebral giant cells. An *ex vivo* study in the pond snail (*Helisoma trivolis*) showed that the spontaneous regeneration of specific neurons was inhibited by serotonin treatments^27^. Serotonin or serotonin reuptake inhibitor treatments also inhibited neurite outgrowth from goldfish retinal explants with a previous crush to the optic nerve^29,30^. So, our results confirm these previous *in vitro* studies and show that endogenous serotonin inhibits axon regeneration after a traumatic injury *in vivo* and in a vertebrate species. The only exception to these results comes from a recent study showing that in the nematode *Caenorhabditis elegans* serotonin promotes axon regeneration^31^. In these animals, non-serotonergic neurons temporarily express tryptophan hydroxylase (the rate-limiting enzyme in serotonin synthesis) in response to axotomy promoting their regeneration^31^. We should take into account that serotonin can signal through a variety of serotonin receptors that activate different secondary pathways. Our results showed that in lampreys the negative effect of endogenous serotonin on the regeneration of descending neurons is caused by the activation of serotonin 1A receptors and a subsequent reduction in cAMP levels. Previous *in vitro* studies in goldfish also showed that the activation of serotonin 1A receptors inhibits neurite outgrowth^29,39^. Accordingly, serotonin 1A agonist treatments reduced neurite outgrowth from retinal explants of goldfish with a prior crush of the optic nerve^29,39^. Interestingly, in *C. elegans* the opposite effect of serotonin was mediated by the activation of serotonin 7 receptors, since mutations of this receptor caused defects in axon regeneration after axotomy^31^. Notably, Alam and co-workers^31^ showed that the positive effect of the activation of this receptor was caused (at least partially) by the stimulation of adenylate cyclase to produce cAMP. Taken together, present and previous results indicate that serotonin is an important regulator of axon regeneration by modulating cAMP levels and that the effects of serotonin will depend on the preferential use of different types of serotonin receptors in different neurons.

Our RNA sequencing study also revealed changes in gene expression in the brainstem associated to serotonin 1A receptor signalling after a complete SCI. Of interest, a treatment with a serotonin 1A receptor antagonist (which increases cAMP levels and promotes axon regeneration) reduced the expression of genes associated to the regulation of axonal guidance. Specifically, Plexin-B1 was one of the most significantly downregulated genes together with the pathways “Signalling by ROBO receptors” and “Regulation of expression of SLITs and ROBOs”. Interestingly, previous work in lampreys has suggested that the expression/activity of some receptors that participate in axonal guidance inhibits axonal regeneration after SCI in identifiable neurons (e.g. UNC5^40–42^). Also, a previous study already suggested a possible involvement of semaphorin/plexin signalling in recovery after SCI in lampreys based on the occurrence of changes in the expression of semaphorins in the spinal cord after the injury^43^. Future studies should functionally test the possible involvement of Slits and semaphorins in the inhibition of axonal regeneration in descending neurons after SCI in lampreys.

Present and previous work in regenerating species shows that serotonin plays different roles in the process of regeneration after SCI. In zebrafish, endogenous serotonin promotes motor neuron production in the spinal cord after a complete SCI by enhancing the proliferation of motor neuron progenitor cells^44^. In turtles, serotonin inhibits the emergence of serotonergic interneurons after SCI by inhibiting a change in neurotransmitter phenotype of non-serotonergic neurons^45^. On the other hand, the production of new-born serotonergic neurons in the spinal cord of zebrafish after SCI is not affected by endogenous serotonin^44^. Here, our results show that in regenerating vertebrates endogenous serotonin also controls axon regeneration of descending neurons after SCI. This indicates that serotonin could also be a target of interest in non-regenerating mammalian models of SCI since it can modulate several aspects of the regenerative process. Even more importantly the effects of serotonin revealed in these studies should be taken into account by those authors performing pharmacological treatments with serotonergic drugs to modulate spinal cord circuits and aiming to promote locomotor recovery after SCI (see ^46^), mainly because these treatments could be affecting the regeneration of new neurons or the re-growth of axotomized axons as shown here.

## Experimental procedures

### Animals

All experiments involving animals were approved by the Bioethics Committee at the University of Santiago de Compostela and the *Consellería do Medio Rural e do Mar* of the *Xunta de Galicia* (license reference JLPV/IId; Spain) and were performed in accordance to European Union and Spanish guidelines on animal care and experimentation. During the experimental procedures, special effort was taken to minimize animal suffering and to reduce the use of animals. Animals were deeply anaesthetized with 0.1% MS-222 (Sigma, St. Louis, MO) in lamprey Ringer solution before all experimental procedures and euthanized by decapitation at the end of the experiments.

Mature and developmentally stable larval sea lampreys, *Petromyzon marinus* L. (n = 176; between 90 and 120 mm in body length, 5 to 7 years of age), were used in the study. Larval lampreys were collected from the river Ulla (Galicia, north-western Spain), with permission from the *Xunta de Galicia*, and maintained in aerated fresh water aquaria at 15°C with a bed of river sediment until their use in experimental procedures. Larval lampreys were randomly distributed between the different experimental groups.

### Spinal cord injury surgical procedures

Larval sea lampreys were assigned to the following experimental groups: control animals without a complete spinal cord transection or animals with a complete spinal cord transection that were analysed 1 wpl, 4 wpl or 11 wpl. Within some of the 4 wpl and 11 wpl groups the animals were assigned to either a vehicle treated control group or to a treatment group. Table 2 summarizes the number of animals assigned to each experimental group. Each experiment was carried out in at least 2 different batches of animals. Complete spinal cord transections were performed as previously described^47^.

Briefly, the spinal cord was exposed from the dorsal midline at the level of the 5^th^ gill by making a longitudinal incision with a scalpel (#11). A complete spinal cord transection was performed with Castroviejo scissors and the spinal cord cut ends were visualized under the stereomicroscope. Then, the animals were kept on ice for 1 hour to allow the wound to air dry. After this hour, the animals were returned to fresh water tanks and each transected animal was examined 24 hours after surgery to confirm that there was no movement caudal to the lesion site. Then, the animals were allowed to recover in fresh water tanks at 19.5 °C and in the dark.

### Drug treatments

The following drugs were used to treat the animals following the complete spinal cord transection: serotonin-hydrochloride (a serotonin analogue that crosses the blood brain barrier; AlfaAesar; Cat#: B21263; MW: 212.68 g/mol), WAY-100,135 (a 5-HT1A antagonist that crosses the blood brain barrier; Sigma; Cat#: W1895; MW: 468.46 g/mol) and db-cAMP (Sigma; Cat#: D0260; MW: 491.37 g/mol). Serotonin-hydrochloride was applied at a concentration of 500 µM in the water were the animals were left after the surgical procedures. Controls were left in fresh water only. Animals were treated with serotonin-hydrochloride for 4 wpl changing the water and the drug twice per week. WAY-100,135 was dissolved in lamprey Ringer’s solution (137 mMNaCl, 2.9 mMKCl, 2.1 mM CaCl2, 2 mM HEPES; pH 7.4) and injected intraperitoneally at a concentration of 1 mM (volume of 25 µl per injection). Vehicle injections served as a control. Animals were treated with WAY-100,135 for 1 or 4 wpl receiving 1 intraperitoneal injection per week. Dd-cAMP was also dissolved in lamprey Ringer’s solution at a concentration of 100 mM, soaked in a small piece of Gelfoam (Pfizer; New York, NY) and placed on top of the site of injury at the time of transection as previously described by other authors^23,24^. Gelfoam soaked in the same volume of lamprey Ringer’s solution served as a control.

We assumed that these drugs also crossed the blood brain barrier in lampreys as in mammals, since the blood brain barrier of lampreys is similar to that of higher vertebrates^48,49^. Serotonin was applied at the same concentration previously used by other authors^46^. Becker and Parker^46^ already reported that this concentration of serotonin affects the swimming behaviour of lesioned and unlesioned animals indicating that this application route also allows access to the CNS. We also observed changes in the swimming behaviour of unlesioned animals in pilot experiments using WAY-100,135 at this concentration (not shown). For db-cAMP application we followed the protocol used by other authors in lampreys in studies in which this treatment promoted axonal regeneration after SCI^23,24^.

### qPCR

Animals with a complete spinal cord transection (4 wpl) and control sham animals without a spinal cord transection (injury of the body wall only) were processed for a qPCR analysis of possible changes in serotonin 1a receptor expression in the whole brainstem. The brainstems of larvae were dissected out and put in RNA*later®* (Ambion Inc., Waltham, MA). RNA extraction was performed using the RNeasy mini kit (Qiagen; Hilden, Germany) with DNase treatment following manufacturer’s instructions. 4 samples were processed in both experimental groups and each sample contained 2 brainstems. RNA quality and quantity were evaluated in a Bioanalyzer (Bonsai Technologies; Madrid, Spain) and in a NanoDrop^®^ ND-1000 spectrophotometer (NanoDrop Technologies Inc; Wilmington, DE), respectively. To determine differential expression between conditions the ΔΔCT method was used with the controls as normalizer, and the Glyceraldehyde 3-phosphate dehydrogenase (GAPDH) as reference gene. Primer sequences for serotonin 1a receptor and gapdh can be found in Supplementary Table 3. Reactions were performed using a qPCR Master Mix Plus SYBR Green I No Rox (Eurogenetec; Liege, Belgium) following the manufacturer instructions and qPCR was carried out on a MX3005P (Agilent Technologies; Santa Clara, CA), including two technical replicates of each sample. Analyses were performed using the MxPro software.

### In situ hybridization

For serotonin 1a receptor *in situ* hybridization, the head of the animals was fixed by immersion in 4% paraformaldehyde (PFA) in 0.05 M Tris-buffered saline (TBS; pH 7.4) for 12 hours. Then, the brains were dissected out, washed and embedded in Neg 50TM (Microm International GmbH, Walldorf, Germany), frozen in liquid nitrogen-cooled isopentane, sectioned on a cryostat in the horizontal plane (14 μm thick) and mounted on Superfrost Plus glass slides (Menzel, Braunschweig, Germany). *In situ* hybridization with a specific riboprobe for the serotonin 1a receptor (GenBank accession number KU314442.1) was conducted as previously described^34,50^. Briefly, brain sections were incubated with the sea lamprey serotonin 1a receptor DIG-labelled probe at 70 °C and treated with RNAse A (Invitrogen, Massachusetts, USA) in the post-hybridization washes. Then, the sections were incubated with a sheep anti-DIG antibody conjugated to alkaline phosphatase (1:2000; Roche, Mannhein, Germany) overnight. Staining was conducted in BM Purple (Roche) at 37°C. *In situ* hybridization experiments were performed in parallel with animals from the different experimental groups (control, 1 wpl and 4 wpl) and the colorimetric reaction was stopped simultaneously for all sections from the different groups of animals.

### Morpholino treatments

The spinal cord was transected at the level of the 5th gill (see surgical procedures), and 1 µl of the morpholinos (0.25 mM in Milli-Q-water) were applied on the rostral stump of the spinal cord. The morpholinos were designed by GeneTools, LLC (Philomath, OR) and included a fluorescein conjugated active translation-blocking serotonin 1a receptor morpholino (5’ - CTGTGATGTTGTGAGCTTCCATCG - 3’) generated against the translation initiation site of the sea lamprey serotonin 1A receptor sequence (Suppl. Fig. 2A) and the fluorescein conjugated GeneTools standard control morpholino (5’-CCTCTTACCTCAGTTACAATTTATA - 3’). During recovery, the morpholinos are retrogradely transported to the cell soma of descending neurons where they can knockdown the expression of the target mRNA (Suppl. Fig. 2B; ^42,51–54^). The GeneTools standard control morpholino has been already used in previous studies using morpholino treatments after SCI in lampreys^51,53^. Animals were allowed to recover for 1, 4 or 11 wpl.

Immunofluorescence experiments were carried out to confirm that the active serotonin 1A receptor morpholino decreases the expression of the sea lamprey receptor. For immunohistochemistry, the brains of larvae were fixed by immersion in 4% PFA in 0.05 M Tris-buffered saline pH 7.4 (TBS) for 4 hours at 4 °C. The samples were then rinsed in TBS cryoprotected with 30% sucrose in TBS, embedded in Tissue Tek (Sakura), frozen in liquid nitrogen-cooled isopentane, and cut serially on a cryostat (14 µm thickness) in transverse planes. Sections were mounted on Superfrost^®^ Plus glass slides (Menzel). The sections were incubated with a rabbit polyclonal anti-serotonin 1A receptor antibody (dilution 1:200; Abcam; Cambridge, UK; Cat#: ab85615; lot GR318082-1; RRID: AB_10696528; immunogen: synthetic peptide corresponding to rat serotonin 1A receptor aa 100-200 conjugated to keyhole limpet haemocyanin) at room temperature overnight. Primary antibodies were diluted in TBS containing 15% normal goat serum and 0.2% Tween as detergent. The sections were rinsed in TBS and incubated for 1 hour at room temperature with a cocktail of Cy3-conjugated goat anti-rabbit (1:200; Millipore; Burlington, MA). Amino acids 100 to 200 of the rat receptor show an 87% correspondence with the same region of the sea lamprey serotonin 1A receptor. Moreover, immunostaining of sea lamprey sections was abolished after pre-adsorption of the anti-serotonin 1A receptor antibody with the synthetic peptide (Suppl. Fig. 2E).

### Behavioural analyses

The behavioural recovery of the animals treated with serotonin-hydrochloride was analysed at 11 wpl based on the study of Ayers^55^ and following the protocol of Hoffman and Parker^56^. This qualitative assessment of locomotor function was made from video recordings of 5 minutes (camera: Panasonic Full-HD HC-V110). The animals were placed in a plastic aquarium (36 ×; 23 ×; 10.5 cm) and swimming activity was initiated by lightly pinching the tail of the animal using a pair of forceps. Locomotor recovery of the animals was categorized in a scale of 1 to 6^55,56^. Animals in stage 5 or 6 correspond to animals in which regeneration of axons through the site of injury has occurred based on activity evoked by stimulation across the lesion site in the isolated spinal cord^56^. Two blinded experimenters independently evaluated each 11 wpl animal. Based on both analyses, a mean value of locomotor recovery was obtained for each animal.

### Retrograde labelling of regenerated descending neurons

At 11 wpl a second complete spinal cord transection was performed 5 mm below the site of the original transection and the retrograde tracer Neurobiotin (NB, 322.8 Da molecular weight; Vector Labs, Burlingame, CA) was applied in the rostral end of the transected spinal cord with the aid of a minute pin (#000). The animals were allowed to recover at 19 °C with appropriate ventilation conditions for 1 week to allow transport of the tracer from the application point to the neuronal soma of descending neurons (the M1, M2, M3, I1, I2, I3, I4, I5, I6, B1, B2, B3, B4, B5, B6, Mth and Mth’ neurons were analysed; Suppl. Fig. 3). Since the original SCI also was a complete spinal cord transection, only neurons whose axons regenerated at least 5 mm below the site if injury were labelled by the tracer. Brains of these larvae were dissected out, and the posterior and cerebrotectal commissures of the brain were cut along the dorsal midline, and the alar plates were deflected laterally and pinned flat to a small strip of Sylgard (Dow Corning Co., USA) and fixed with 4% PFA in TBS for 4 hours at 4 °C. After washes in TBS, the brains were incubated at 4 °C with Avidin D-FITC conjugated (Vector; Cat#: A-2001; 1:500) diluted in TBS containing 0.3% Triton X-100 for 2 days to reveal the presence of Neurobiotin. Brains were rinsed in TBS and distilled water and mounted with Mowiol.

### Assay for the quantification of cAMP concentration

cAMP concentration in the brainstem of animals treated with WAY-100,135 was analysed 1 wpl using a quantitative cAMP competitive ELISA kit (Invitrogen, Waltham, MA, USA; Cat#EMSCAMPL) following the manufacturer’s instructions. This assay is based on the competition between cAMP in the standard or sample and alkaline phosphatase conjugated cAMP for a limited amount of cAMP monoclonal antibody bound to an anti-rabbit IgG precoated 96-well plate. The brains of larvae were dissected out 1 hour after the second WAY-100,135 or control intraperitoneal injections and immediately homogenized in 0.1 M HCl on ice.

### RNA sequencing

Animals treated during 4 wpl with WAY-100,135 and control vehicle treated animals were processed for an RNA sequencing analysis of the whole brainstem. The brainstems of larvae were dissected out and immediately put in RNA*later^®^* (Ambion Inc.). RNA extraction was performed using the RNeasy mini kit (Qiagen) with DNase treatment following manufacturer’s instructions. RNA quality and quantity were evaluated in a Bioanalyzer (Bonsai Technologies) and in a NanoDrop^®^ ND-1000 spectrophotometer (NanoDrop^®^ Technologies Inc), respectively. Three WAY-100,123 and three control samples (each sample containing 4 brainstems) were barcoded and prepared for sequencing by the Wellcome Trust Centre for Human Genetics (Oxford, UK) using standard protocols. Sequencing was conducted on an Illumina HiSeq 2000 as 100 bp paired-end reads. Raw sequencing has been deposited in NCBI’s Short Read Archive (SRA) under BioProject accession PRJNA472778. The quality of the sequencing output was assessed using FastQC v.0.11.5. (http://www.bioinformatics.babraham.ac.uk/projects/fastqc/). Quality filtering and removal of residual adaptor sequences was conducted on read pairs using Trimmomatic v.0.35^57^. Specifically, residual Illumina specific adaptors were clipped from the reads, leading and trailing bases with a Phred score less than 20 were removed and the read trimmed if a sliding window average Phred score over four bases was less than 20. Only reads where both pairs had a length greater than 36 bp post-filtering were retained. A *de novo* transcriptome was assembled using Trinity v.2.4.0^58^ with default settings. Gene expression was estimated using Kallisto v.0.43.1^59^ and statistical analyses related to differential expression were performed using R v.3.4.3 (R Core Team 2014). Gene count data were used to estimate differential gene expression using the Bioconductor package DESeq2 v.3.4^60^. The Benjamini-Hochberg false discovery rate (FDR) was applied, and transcripts with corrected p-values < 0.05 were considered differentially expressed genes. Heatmaps were drawn using the R package “gplots” v.3.01.1^61^. Pathway analyses were performed using Reactome^38^, converting all non-human gene identifiers to their human equivalents.

### Imaging and quantifications

The percentage of neurons with regenerated axons (labelled by the Neurobiotin tracer) respect to the total number of analysed neurons (see Table 1) was calculated for each type of identifiable descending neuron using an Olympus microscope. The percentage of neurons with regenerated axons respected to the total number of analysed neurons in each animal was also calculated and these data was used for statistical analyses. The experimenter was blinded during quantifications. For the figures, images were taken with a spectral confocal microscope (model TCS-SP2; Leica).

An Olympus photomicroscope (AX-70; Provis) with a 20x Apochromatic 0.75 lens and equipped with a colour digital camera (Olympus DP70, Tokyo, Japan) was used to acquire images of brain sections from the *in situ* hybridization experiments. Images were always acquired with the same microscope and software settings. The quantification of the level of serotonin 1a receptor positive signal in identifiable descending neurons was performed as previously described^54^. First, we established the intensity rank of positive colorimetric *in situ* signal. For this, we analysed 10 random images from different descending neurons of control and lesioned animals. The “histogram” function in Image J shows the number of pixels in each image in a range of intensity from 0 to 255. With these images, we compared the intensity values in regions with clear *in situ* signal and the intensity values in regions without *in situ* signal. Based on this, we established a value of 179 as the lower limit to consider the colorimetric *in situ* signal as positive. Then the number of pixels of positive *in situ* signal was quantified for each section of each identified descending neuron. In horizontal brain sections, the identification of some of the specific descending cells becomes more difficult than in whole-mounts. Thus, only the cells that were unequivocally identified in at least two different sections were included in the quantifications (the M1, M2, M3, I1 and I3 neurons; see Suppl. Fig. 3). Then, we calculated the average number of positive pixels per section for each individual neuron (see Table 1) and this data was used for statistical analyses. The experimenter was blinded during quantifications.

For the quantification of changes in serotonin 1A receptor immunoreactivity after morpholino application, 3 14-µm transverse sections of the medial reticular nucleus of the rhombencephalon were analysed in each animal. 1 out of every 2 consecutive sections in the next 6 sections caudally to the last section where the Mauthner neuron was observed were analysed in each animal. The mean fluorescence intensity of each section was calculated using Image J and then the mean fluorescence intensity per section in each animal was used for the statistical analyses.

After quantifications, contrast and brightness were minimally adjusted with Adobe Photoshop CS6 (Adobe Systems, San José, CA, USA). Figure plates and lettering were generated using Adobe Photoshop CS6 (Adobe Systems). Schematic drawing was generated using CorelDraw Graphics Suite 2017.

### Statistical analyses

Statistical analysis was carried out using Prism 6 (GraphPad software, La Jolla, CA). Data were presented as mean ± S.E.M. Normality of the data was determined by the Kolmogorov-Smirnov test or the D’Agostino-Pearson omnibus test. The data from the db-cAMP treatments and the *in situ* hybridization data that were normally distributed were analysed by a one-way ANOVA. Post-hoc Dunnetts multiple comparison tests were used to compare pairs of data. *In situ* hybridization data that were not normally distributed were analysed by a Kruskal-Wallis test and post-hoc Dunn’s multiple comparisons test. A Student’s t-test was used to determine significant differences between conditions in the qPCR analysis. The results of control versus treatment groups were analysed by a Student’s t-test (normally distributed data) or Mann Whitney U test (non-normally distributed data). The significance level was set at 0.05. In the figures, significance values were represented by a different number of asterisks in the graphs: 1 asterisk (*p* value between 0.01 and 0.05) and 2 asterisks (*p* value between 0.001 and 0.01). Exact *p* values are given in the text.

## Author Contributions

M.C.R. and A.B.-I. conceived and supervised the study. D.S.-C. carried out all the experimental work (with the exception of the qPCR and RNA-seq experiments, which were performed by D.R. and L.S.). D.S.-C., D.R., M.C.R. and A.B.-I. carried out the analysis and interpretation of experimental data. D.S.-C. and D.R. prepared the figures and tables. A.B.-I. wrote the manuscript with help from the other authors.

## Acknowledgments

Grant sponsors: Spanish Ministry of Economy and Competitiveness and the European Regional Development Fund 2007–2013 (Grant number: BFU2014-56300-P) and Xunta de Galicia (Grant number: GPC2014/030). A.B.-I. was supported by a grant from the *Xunta de Galicia* (Grant number: 2016-PG008) and a grant from the crowdfunding platform *Precipita* (*FECYT*; Spanish Ministry of Economy and Competitiveness; grant number 2017-CP081). The authors would like to acknowledge the following individual donors of the crowdfunding campaign in *Precipita*: Blanca Fernández, Emilio Río, Guillermo Vivar, Pablo Pérez, Jorge Férnandez, Ignacio Valiño, Pago de los Centenarios, Eva Candal, María del Pilar Balsa, Jorge Faraldo, Isabel Rodríguez-Moldes, José Manuel López, Juan José Pita, María E. Cameán, Jesús Torres, José Pumares, Verónica Rodríguez, Sara López, Tania Villares Balsa, Rocío Lizcano, José García, Ana M. Cereijo, María Pardo, Nerea Santamaría, Carolina Hernández, Jesús López and María Maneiro. The authors thank the staff of Ximonde Biological Station for providing lampreys used in this study, and the Microscopy Service (University of Santiago de Compostela) and Dr. Mercedes Rivas Cascallar for confocal microscope facilities and help.

## Competing interests

The authors declare that there are no competing interests.

## Supplementary Figures

**Supplementary Figure 1.** Boxplot showing the expression of serotonin 1a receptor in the brainstem of control samples or samples treated with WAY-100,135. Log2 fold change values have been normalized to the mean of the controls. The means of the two groups were not significantly different (*p* = 0.71).

**Supplementary Figure 2.** Serotonin 1A receptor morpholino. **A:** Partial sequence of the sea lamprey 5-ht1a gene with exon sequence in red and 5’ untranslated sequence in black. The target sequence of the 5-HT1A MO is between square brackets. The ATG initiation codon is between parenthesis. **B:** Photomicrograph of a whole-mounted brain showing the presence of the 5-HT1A MO in identifiable neurons 1week after MO application. **C:** Photomicrographs of transverse sections of the brainstem showing 5-HT1A immunoreactivity in control and 5-HT1A MO treated animals. Note the decreased immunoreactivity in reticulospinal neurons of 5-HT1A MO treated animals. **D:** Graph showing significant changes (asterisks) in the mean fluorescence intensity of 5-HT1A immunoreactivity per section in reticulospinal nuclei of the brainstem after the 5-HT1A MO treatment. **E:** Photomicrographs of transverse sections of the brainstem showing 5-HT1A immunoreactivity in control and in anti-5-HT1A antibody pre-adsorption experiments.

**Supplementary Figure 3.** Schematic drawing of a dorsal view of the sea lamprey brainstem showing the location of identifiable descending neurons (modified from Sobrido-Cameán and Barreiro-Iglesias, 2018). Abbreviations: M: mesencephalon; R: rhombencephalon; SC: spinal cord. Rostral is up.

**Supplementary Tables**

**Supplementary Table 1.** Differentially expressed genes between control and WAY-100,135 treated samples.

List of genes showing significant differential expression (FDR p-value < 0.05) between control samples and samples treated with WAY-100,135. Gene expression level (average normalized count values), log2 fold change (positive values represent up-regulation in response to WAY-100,135 treatment), FDR corrected p-value, Uniprot ID of the most significant blast hit, blast alignment length, blast alignment identity percentage, blast alignment E value, protein name of the most significant blast hit, gene symbol of the most significant blast hit and species of the most significant blast hit are shown for every differentially expressed gene.

**Supplementary Table 2.** Results of the Reactome pathway analysis. List of Reactome pathways showing significant enrichment (FDR p-value < 0.05) for the list of differentially expressed genes between control and WAY-100,135 treated samples. Pathway name, total genes assigned to the pathway, total genes present in the pathway and FDR corrected p-value are shown for every enriched Reactome pathway.

**Supplementary Table 3.** Primer sequences for gapdh and 5-ht1a in qPCR experiments.

